# The FABRIC Cancer Portal: A Ranked Catalogue of Gene Selection in Tumors over the Human Coding Genome

**DOI:** 10.1101/2020.11.26.399543

**Authors:** Guy Kelman, Nadav Brandes, Michal Linial

## Abstract

We present The FABRIC Cancer Portal, a comprehensive catalogue of gene selection in cancer covering the entire human coding genome (~18,000 protein-coding genes) across 33 cancer types and pan-cancer. Genes in the portal are ranked according to the strength of evidence for selection in tumor, based on rigorous and robust statistics. Gene selection is quantified by combining genomic data with rich proteomic annotations. The portal includes a selected set of cross-references to the most relevant resources providing genomic, proteomic and cancer-related information. We showcase the portal with known and overlooked cancer genes, demonstrating the usefulness of the portal via its simple visual interface that allows to pivot between gene-centric and cancer-type views. The portal is available at fabric-cancer.huji.ac.il.

## Introduction

Cancer gene catalogues, annotating accumulated knowledge and evidence about the roles of genes in tumors, are an indispensable tool in cancer research and precision therapy [1, 2]. One of the main goals of such catalogues is to highlight potential cancer drivers [3, 4]. However, lists of candidate cancer drivers are typically limited to a few hundreds of high-confidence genes, without any information about the rest of the coding genome (e.g. 719 genes in Census (V86) [3], 299 genes in [4]). Moreover, gene annotations in driver catalogues are typically binary, marking whether each gene is a potential cancer driver or not, without expressing ranking, effect size or confidence level for candidate genes. Finally, as the primary evidence used to implicate genes in cancer within existing catalogues is mutation abundance, they fail to capture low-recurrence driver genes.

### Outline

We recently developed FABRIC, a new method for detecting genes involved in cancer [5] (Fig. 1). Unlike other computational methods, which mostly consider the recurrence of mutations, FABRIC extracts signal from the molecular functional effects of mutations altering protein-coding genes. In other words, instead of looking for genes with high rates of mutations, FABRIC detects genes with mutations that are more damaging than would be expected at random, regardless of their number. The framework is therefore complementary to most other methods in finding genes under positive selection. We used FABRIC to assess and quantify the selection of the entire human proteome (~18,000 protein-coding genes) across 33 cancer types and pan-cancer, by analyzing ~3M somatic mutations in ~10,000 cancer patients from the TCGA cohort [6]. A full description of that analysis is described in our original work [5]. FABRIC’s summary statistics across the entire human coding genome with respect to the 33 cancer types and pan-cancer provide a wealth of information on the potential roles of human genes in tumors.

**Fig. 1:**
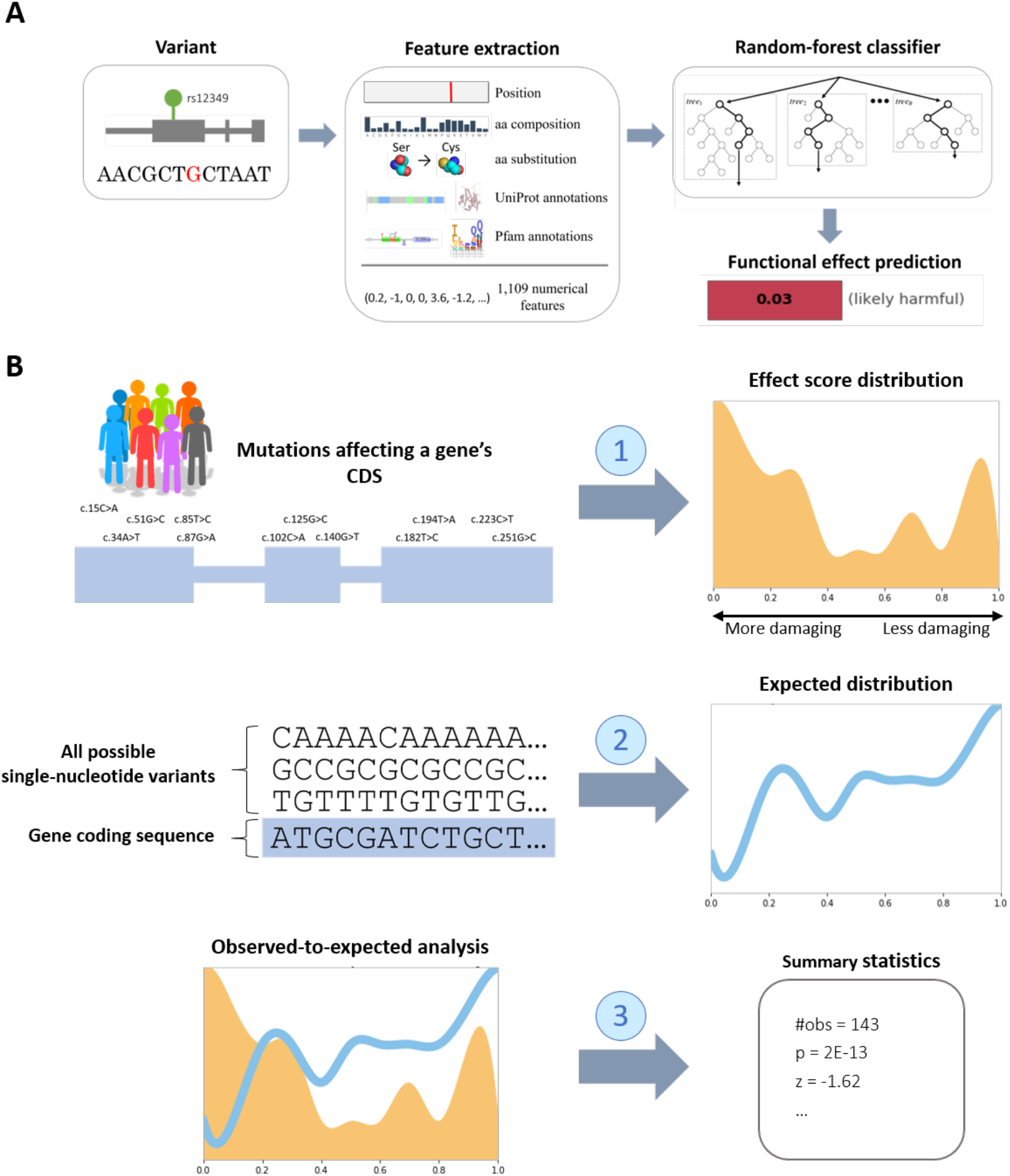
The FABRIC framework. FABRIC (Functional Alteration Bias Recovery In Coding-regions) is a framework for detecting and quantifying gene selection in coding regions. In the context of cancer, it can analyze somatic mutations to identify protein-coding genes under positive selection, which are suspect driver genes. (**A**) FABRIC uses an underlying machine learning model (called FIRM) to estimate the functional damage of genetic variants in coding regions. FIRM extracts a large set of proteomic features from a variety of sources (including UniProt and Pfam annotations). Based on these features, it assigns each mutation an effect score reflecting the probability of the protein to retain its function given the mutation. (**B**) The FABRIC framework consists of three steps. To assess the selection of a gene, FABRIC first collects all single-nucleotide mutations within its coding region (from a variety of samples) and uses FIRM to assign them functional effect scores. Second, FIRM is also used to record the effect scores of all possible single-nucleotide variants within the same gene, in order to construct a background model for the distribution of effect scores that would be expected at random. FABRIC’s background model takes into account the number of observed mutations and their nucleotide substitution distribution. In the third and final step, the effect score distribution of the observed mutations is compared against the distribution of the expected effect scores to obtain the summary statistics for the analyzed gene. These summary statistics describe the significance (p-value) and strength (z-value) of the gene’s selection, namely to what extent the observed mutations are more (or less) damaging than would be expected by the same number of random mutations.

Altogether, FABRIC recovers 593 protein-coding genes exhibiting significant positive selection in pan-cancer, and only 6 genes showing signs of negative selection (Fig. 2). In many cases, FABRIC recovers well-known cancer driver genes such as TP53, APC and KRAS. However, it doesn’t always converge with other methods for detecting caner drivers, especially those based on mutation rates. Unlike most methods, FABRIC considers the functional effects of mutations rather than their number. For example, the FAT4 gene is a tumor suppressor gene playing a role in numerous cancers [7] (also cf Fig. 2, a high observed effect overruled by the background model). Despite its high mutation rate (with 1,893 single-nucleotide somatic mutations in the coding region of the gene across the 33 TCGA cancer types), it is not significant by FABRIC (FDR q-value = 0.11). On the other hand, FABRIC is able to recover hundreds of overlooked genes [5] (e.g. MICU3, FDR q-value = 4E-7).

**Fig. 2:**
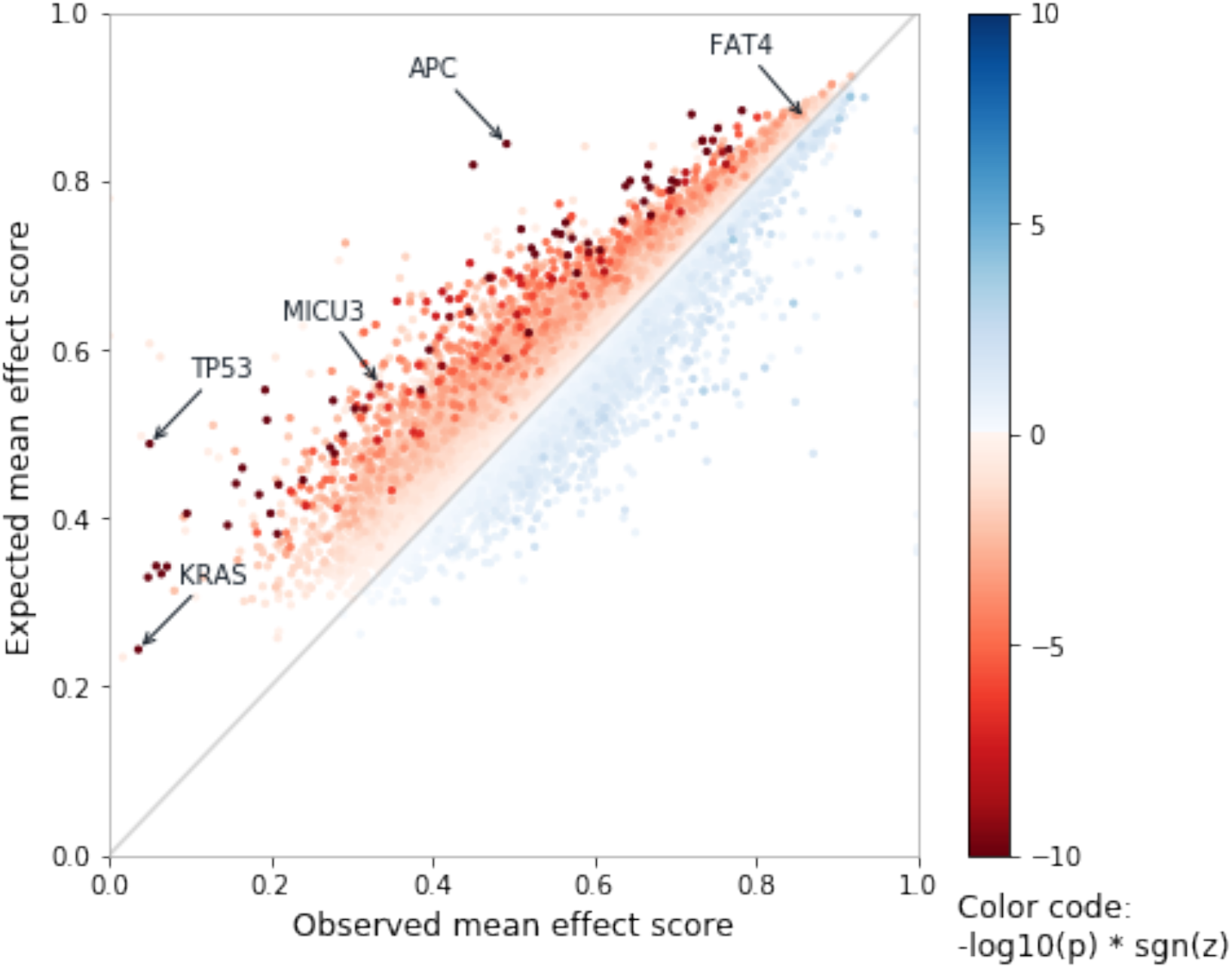
FABRIC summary statistics in pan-cancer across the human coding genome. The observed and expected mean effect scores assigned by FABRIC to each of the ~18,000 analyzed protein-coding genes in pan-cancer. The x-axis marks the average effect score across all somatic mutations observed in a gene, with lower scores indicating more damaging mutations. The y-axis marks the mean effect score that would be expected for the same number of mutations, as calculated by FABRIC’s background model. Genes are colored by their significance (log-scaled p-values). Genes under positive selection (i.e. affected by mutations more damaging than would be expected at random, indicated by negative z-values) are colored red, while in blue are genes under negative selection (i.e. with mutations less damaging than would be expected at random, indicated by positive z-values).

Here we present The FABRIC Cancer Portal, available at: fabric-cancer.huji.ac.il. The goal of the new portal is to make the rich cancer results of FABRIC widely available via a simple web resource that enables quantitative and comparative analyses. In addition to easy access to downloadable data, the portal also provides quick cross-references to other genomic, proteomic and cancer resources (Table 1).

**Table 1:**
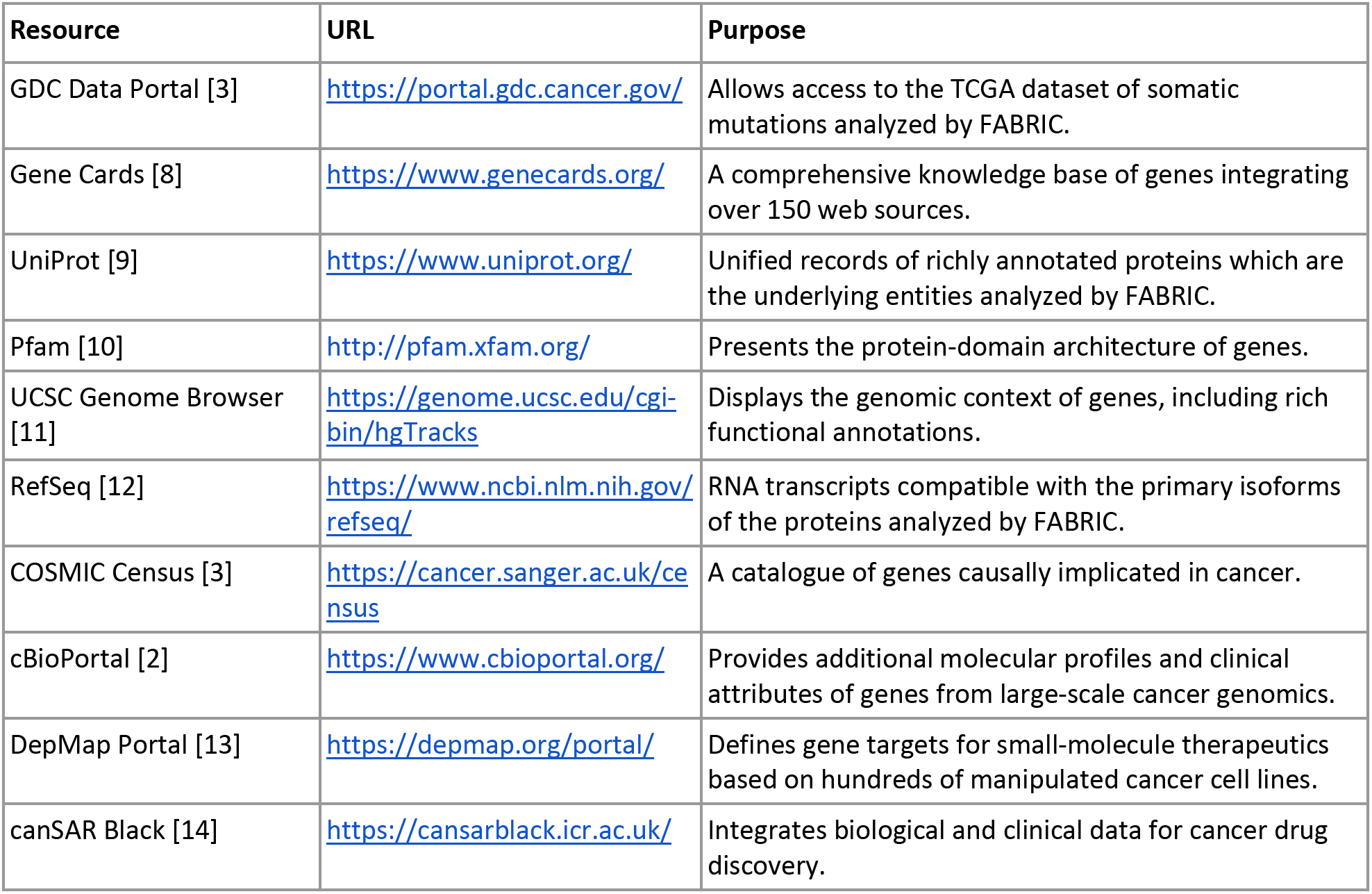
Cross-references in The FABRIC Cancer Portal.

### Showcase

To illustrate the usability of The FABRIC Cancer Portal as a navigation and discovery tool, we present a showcase starting with APC (*Adenomatosis Polyposis Coli)*, a known cancer driver gene (Fig. 3). APC is a classic tumor suppressor gene playing a central role in cell cycle and migration through the Wnt signaling pathway. The role of APC in cell division and differentiation is mediated by its direct binding to β-catenin [15].

**Fig. 3:**
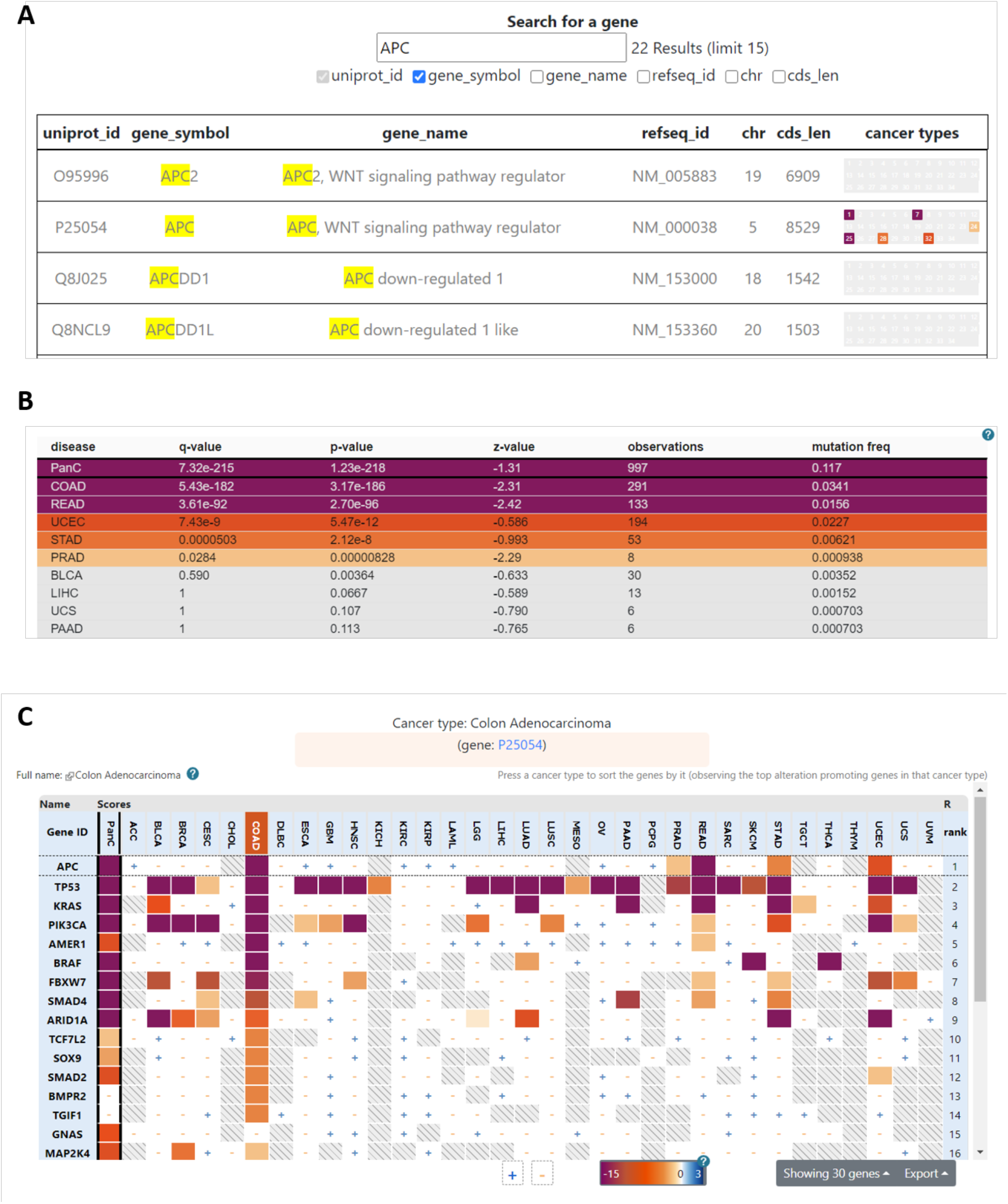
Searching a gene and viewing selection patterns across cancer types (screenshots) (**A**) Typing “APC” in the portal’s search page (fabric-cancer.huji.ac.il/g/search). (**B**) Selecting the APC gene links to the gene’s page (fabric-cancer.huji.ac.il/g/gene/APC), with a table showing FABRIC’s summary statistics for APC across 33 cancer types and pan-cancer. (**C**) Choosing the COAD (colon adenocarcinoma) cancer type within that table switches to the primary heatmap, centered around the intersection of COAD and APC (fabric-cancer.huji.ac.il/g/disease/COAD?focus=P25054). The heatmap is color-coded by the significance of gene and cancer-type pairs.

The FABRIC Cancer Portal allows us to observe the significance (p-value and FDR q-value) and effect size (z-value) of a gene’s selection across cancer types. On top of the 33 individual cancer types, it also jointly examines the somatic mutations from all cancer types aggregated, an analysis referred to as pan-cancer. We find that APC is specific to 5 cancer types (Fig. 3B), all of them are related to the digestive tract. In particular, the gene is under very significant and very strong positive selection in colorectal cancers, colon adenocarcinoma (COAD; q = 5E-182, z = −2.31) and rectum adenocarcinoma (READ; q = 3E-96, z = −2.42), indicating that APC is a likely driver in these cancers. To a lesser extent, it also exhibits excess functional damage in uterus cancer (UCEC; q = 7E-9, z = −0.59), stomach cancer (STAD; q = 5E-5, z = −0.99) and prostate cancer (PRAD; q = 0.03, z = −2.29). In addition to the listed cancer types, we also observe APC’s strong pan-cancer signal (q = 7E-215, z = −1.31).

From within APC’s gene page we can choose the COAD cancer type and navigate to the portal’s primary heatmap. This heatmap visualizes the degree of selection across genes and cancer types, centered around the intersection of COAD (colon adenocarcinoma) and APC (Fig. 3C). Having selected that particular cancer type, the heatmap ranks the genes according to their significance in COAD. We observe that APC is in fact ranked first in this cancer type. The next three genes, TP53, KRAS and PIK3CA (Fig. 3C), are also known cancer driver genes, which, as expected, dominate across many other cancer types and the pan-cancer analysis. The gene ranked fifth, AMER1, shows a different pattern. It exhibits a selection pattern that is almost exclusive to colon adenocarcinoma (COAD; q = 5E-18), with milder significance in pan-cancer (PanC; q = 3E-9) and rectum adenocarcinoma (READ; q = 2E-5). Based on this selection pattern, we can infer that the role of AMER1 in cancer is restricted to colorectal cancer.

Selecting AMER1 in the heatmap will navigate to the gene page of AMER1 (Fig. 4A). From the information on the page, we can infer that AMER1 is tightly related to APC. The first indication is the gene’s full name: *APC membrane recruitment protein 1*. Further inspection of AMER1 can be carried out by following the available cross-references (Table 1). For example, pressing the COSMIC cross-reference logo links to the AMER1 gene page on COSMIC Census (Fig. 4B). There, we find that AMER1 is indeed annotated as a tumor suppressor. Like APC, it promotes β-catenin ubiquitination and degradation, playing a role in the inhibition of cell growth by induction of G1/G0 arrest [16]. The protein-family domain organization of AMER1 according to Pfam (Fig. 4A) highlights numerous segments of low complexity which often signify unstructured regions. In addition, it contains two structural WTX domains (Wilms’ tumor suppressor X chromosome). This domain architecture is shared among all human paralogs of AMER1, namely AMER2 and AMER3. Further information from GeneCards covers additional aspects of the gene, including its clinical implications, regulation, expression and protein-protein interactions. Overall, we receive confirmation that AMER1 binds directly to APC and, as a result, acts to inhibit Wnt signaling by inducing β-catenin degradation. From canSAR and DepMap we observe that no approved drug targeting AMER1 is under clinical investigation for the treatment of bowel cancers.

**Fig. 4:**
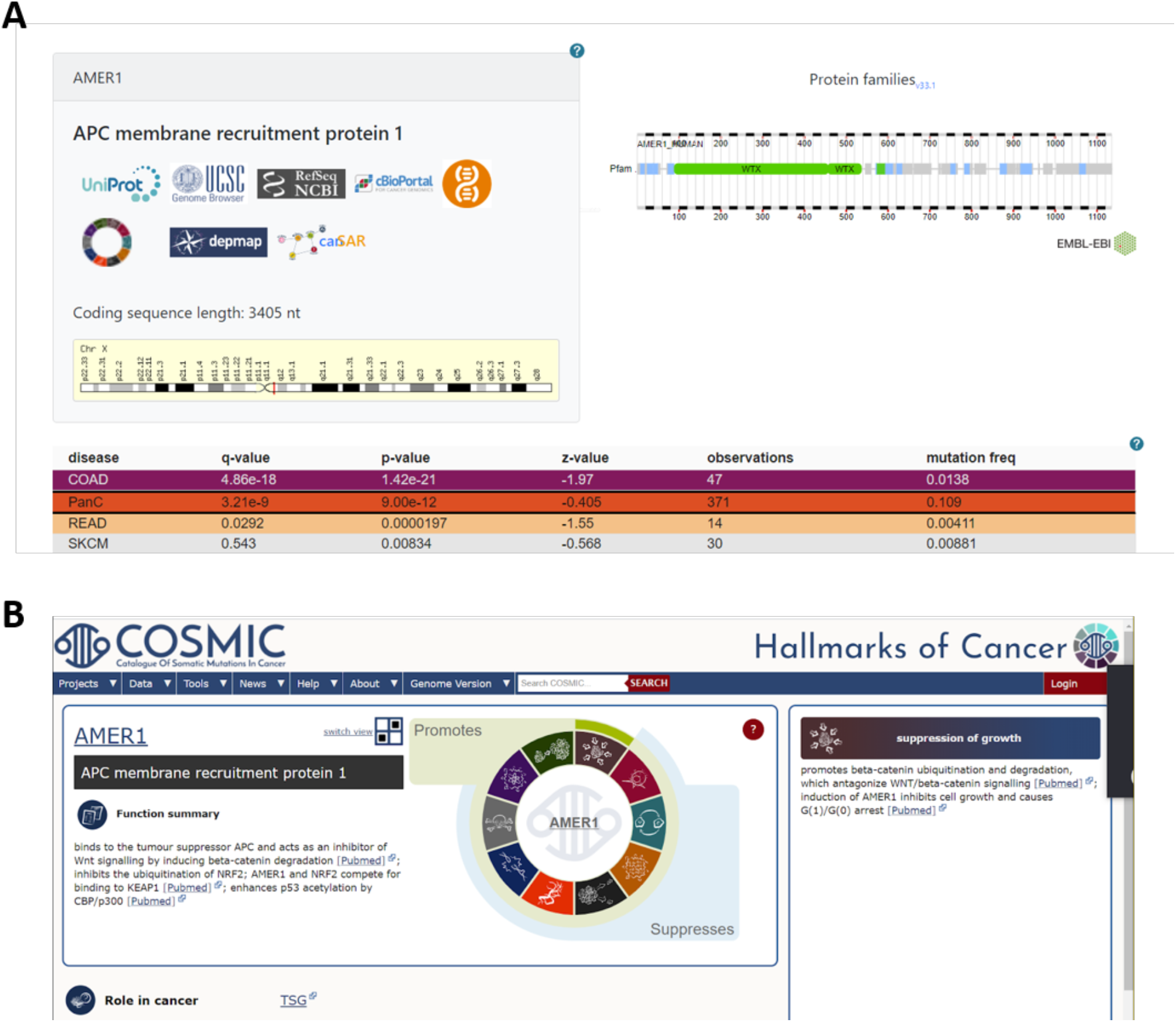
The gene page and COSMIC cross-reference (screenshots) (**A**) The AMER1 gene page includes a table at the bottom with the summary statistics of FABRIC across cancer types and, above it, the protein domain architecture of the gene according to Pfam (upper-right panel) and cross references to other prominent proteomic and cancer web resources with information about the gene (upper-left panel). The gene’s position overlaid on the chromatin G-bands of chromosome X appears on this panel too. (**B**) Pressing the COSMIC Hallmarks icon from the cross-references section links to the gene’s page on COSMIC.

In addition to known cancer driver genes such as APC and AMER1, The FABRIC Cancer Portal also lists hundreds of previously overlooked genes that are positively selected in cancer [5]. One of these genes is MICU3 (*mitochondrial calcium uptake family member 3*), a gene observed to be mutationally active in uterine corpus endometrial carcinoma (UCEC). A possible entry point to UCEC (and other cancer types) is through the body-parts view in the Cancer Types page on the portal (Fig. 5A). Choosing uterus as a primary site shows that MICU3 is ranked 41st in UCEC on the heatmap.

**Fig. 5:**
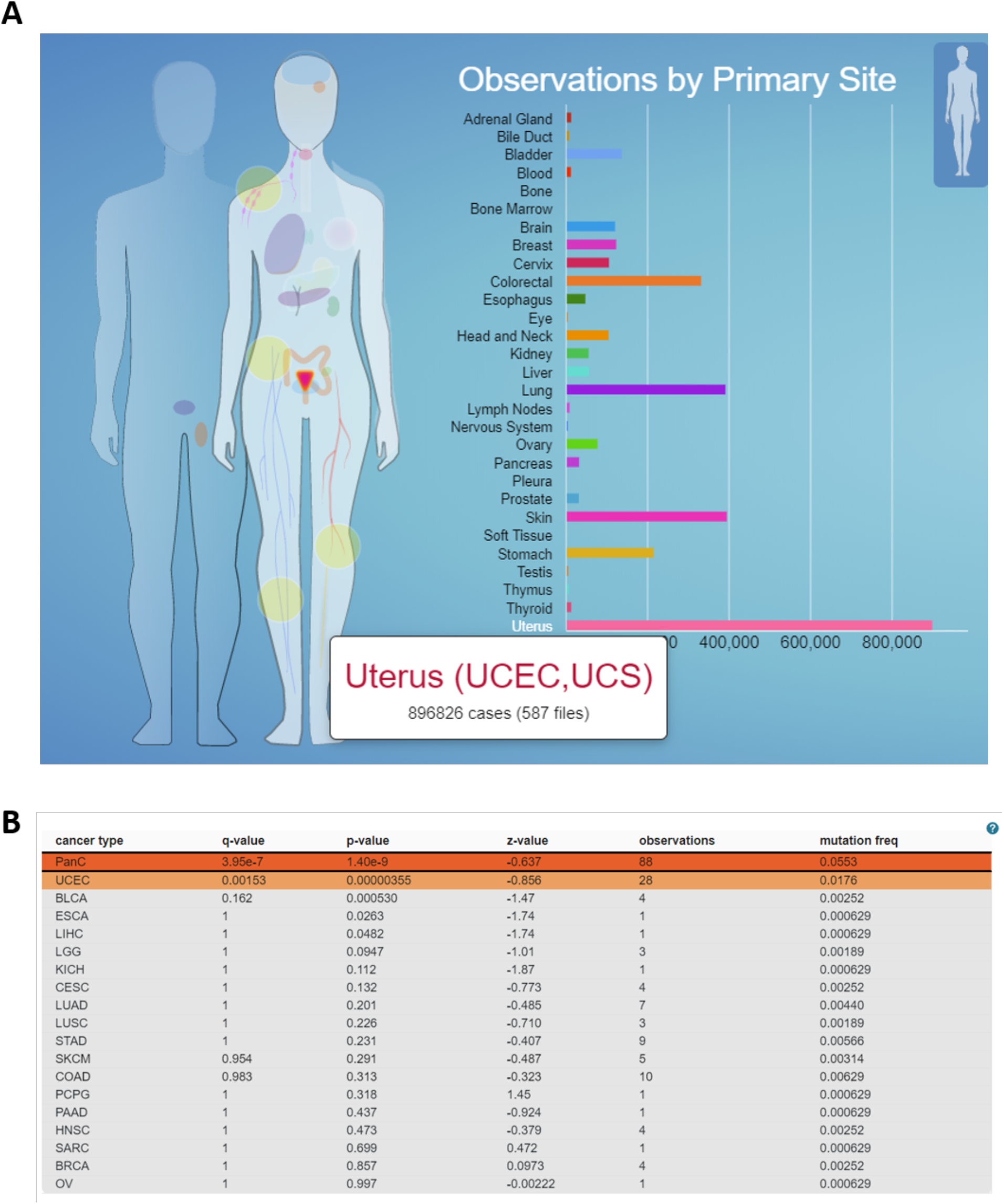
Body-parts view and the MICU3 gene (screenshots) (**A**) The body-parts view in The FABRIC Cancer Portal (adapted from the GDC Portal). For each primary site, the numbers of samples and mutations are provided. In the uterus (UCEC & UCS), for example, TCGA lists 896,826 somatic mutations from 587 samples. (**B**) FABRIC’s summary statistics from the gene page of MICU3 (*mitochondrial calcium uptake family member* 3), a gene overlooked by other cancer gene catalogues.

Notably, MICU3 is not listed as a canonical cancer driver gene by any of the major cancer catalogues [3, 4]. According to FABRIC, however, it is highly significant (Fig. 5B). The positive selection of MICU3 is only significant in uterine corpus endometrial carcinoma (UCEC; q = 0.0015). Nonetheless, the pan-cancer analysis is able to pick an even stronger signal (PanC; q = 4E-7), suggesting that the gene is mildly involved in other cancers as well.

Inspection of the biological and cellular roles of MICU3 through gene knowledgebases such as GeneCards and UniProt brings up relevant facts about the gene. MICU3 is a stimulator of mitochondrial Ca2+ uptake through the mitochondrial calcium uniporter (MCU) complex [17]. Numerous evidence links dysfunctional MCU to cancer [18], and a recent cell-line study suggests that mutations in MICU3 may lead to cell invasion by altering Ca2+ dynamics [19], pointing to MICU3 as a likely cancer driver gene candidate. The strong positive selection of the gene in uterine corpus endometrial carcinoma (and potentially other cancer types) as quantified by FABRIC provides a very strong case that MICU3 is indeed involved in cancer.

The exemplified case studies of APC, AMER1 and MICU3 demonstrate the power of The FABRIC Cancer Portal as an interactive navigation tool across genes and cancer types promoting new discovery.

## Conclusion

We have introduced The FABRIC Cancer Portal, a new resource for exploring the selection patterns of human coding genes in cancer. We have showcased the use of the portal as a discovery tool highlighting overlooked genes and providing new angles on established cancer driver genes.

FABRIC minimizes modeling assumptions and compares each gene against its own background, making it highly robust to false discoveries. The model combines genomic and proteomic data, incorporating updated high-quality proteomic annotations (including phosphorylation and other post-translational modifications, protein domains and secondary structure annotations). Importantly, FABRIC only analyzes coding genes, while ignoring expression data and regulatory elements of the genome (including splicing patterns).

The FABRIC Cancer Portal provides a comprehensive catalogue of the entire human coding genome across all 33 TCGA cancer types. On top of specific cancer types, the portal also provides a pan-cancer view that can capture more subtle signals that might be missed from individual cancer types due to insufficient number of observations. This is especially important in the detection of lowly recurrent genes. As demonstrated, the portal can easily pivot between gene-centric and cancer-type views. Within each cancer type, FABRIC provides a clear ranking of genes by significance. From the gene view, the portal cross-references to a handful of selected external resources providing high-quality information from genomic, proteomic and clinical perspectives. The entire portal data and the results of specific queries are easily downloadable.

## Accessibility

The FABRIC Cancer Portal is available at fabric-cancer.huji.ac.il.

To learn more about its applicability, we invite the reader to browse through the tutorial (fabric-cancer.huji.ac.il/#tutorial-head) and FAQ (fabric-cancer.huji.ac.il/g/FAQ) sections of the portal.

## Acknowledgements

The results in The FABRIC Cancer Portal are based upon data generated by the TCGA Research Network: https://www.cancer.gov/tcga.

## Notes

Conflict of interest: The authors declare no potential conflicts of interest.

http://fabric-cancer.huji.ac.il

### Competing Interest Statement

The authors have declared no competing interest.

### Summary of Updates

The last word of the textual abstract contains a link to the website, but currently it does not show as a clickable link. We made a change to the last word in the abstract in the hope that adding 'http://' in front of it will give the desired outcome

